# Deep6mA: a deep learning framework for exploring similar patterns in DNA N6-methyladenine sites across different species

**DOI:** 10.1101/2019.12.28.889824

**Authors:** Zutan Li, Hangjin Jiang, Lingpeng Kong, Yuanyuan Chen, Liangyun Zhang, Cong Pian

## Abstract

N6-methyladenin(6mA) is an important DNA modification form associated with a wide range of biological processes. Identifying accurately 6mA sites on a genomic scale is crucial for understanding of 6mA’s biological functions. In this paper, we developed, without requiring any prior knowledge of 6mA and manually crafted sequence features, a deep learning framework named Deep6mA to identify DNA 6mA sites, and its performance is superior to other DNA 6mA prediction tools. Specifically, the 5-fold cross-validation on a benchmark dataset of rice gives the sensitivity and specificity of Deep6mA as 92.96% and 95.06%, respectively, and the overall prediction accuracy is 94%. Importantly, we find that the sequences with 6mA sites share similar patterns across different species. The model trained with rice data predicts well the 6mA sites of other three species: *Arabidopsis thaliana*, *Fragaria vesca*, and *Rosa chinensis*, with a prediction accuracy over 90%. In addition, we find that (1) 6mA tends to occur at GAGG motifs, which means the sequence near the 6mA site may be conservative; (2) 6mA is enriched in the TATA box of the promoter, which may be the main source of its regulating downstream gene expression.

## INTRODUCTION

DNA methylation modifications such as such as N4-methylcytosine (4mC), N6-methyladenine (6mA), and 5-methylcytosine (5mC) play important roles in epigenetic regulation of gene expression without altering the sequence, and it is widely distributed in the genome of different species (1). DNA N6-methyladenine (6mA) refers to the methylation of the 6th position of purine ring of adenine, which is one of the most abundant DNA modifications found in eukaryotes and prokaryotes (2). Researches on 6mA shown that 6mA plays important roles in DNA repair (3,4), DNA replication (5), regulating gene transcription (6) and gene expression regulation (7). Although 6mA sites are not uniformly distributed across the genome and they may be affected by environmental factors (8), the methylation protection is a genetic state, and 6mA in prokaryotes and eukaryotes shows similar characteristics (9). It is indispensable to study the distribution of DNA 6mA on the genome for a deeper understanding its epigenetic modification process.

Due to recent advances in high-throughput sequencing technologies, various experimental techniques were reported to promote the study of 6mA distribution and its potential function in genome of eukaryotes and prokaryotes. For example, Pormraning et al. (2009) applied sequencing of methylated DNA immunoprecipitation technique to analyze genome-wide DNA methylation in eukaryotes. Krais et al. (2010) reported that the capillary electrophoresis with laser-induced fluorescence can be used to detect global adenine methylation in DNA. Meanwhile, Flusberg et al. (2010) used a method based on single-molecule, real-time sequencing technique to detect DNA methyladenine directly. Greer et al. (2015) used the ultra-high-performance liquid chromatography and the mass spectrometry to discover DNA 6mA sites. These methods advanced the research of 6mA. By using 6mA-IP-Seq, Fu et al. (2015) found that 84% of 6mA modification exit in Chlamydomonas genes. M. J. Koziol et al. (2016) identified the 6mA modification in vertebrates by using HPLC, blots, and sequencing of methylated DNA Immunoprecipitation (MeDIP-seq). Zhou et al. (2018) found through 6mA immunoprecipitation, mass spectrometry, and single molecule realtime that 0.2% of adenines in the rice genome are 6mA methylated and GAGG-rich sequences are the most significantly enriched for 6mA.

To date, tools were developed to predict 6mA methylation modification. For instance, Chen et al. (2019) proposed a method called i6mA-Pred to identify DNA 6mA sites based on the support vector (SVM) with 164 chemical features of nucleotides and position-specific nucleotide frequencies. The i6mA-Pred shows a good classification performance in rice 6mA data. However, it does not fully capture the information between nearby nucleotides. To address this weakness, Pian et al. (2019) proposed to use MM-6mAPred, a first-order Markov model, to predict 6mA sites. They found that there is a significant difference in the transition probabilities between the neighboring nucleotides in 6mA sequences and non-6mA sequences. Compared with i6mA-Pred, MM-6mAPred has a better performance in terms of prediction accuracy.

The above two prediction models were trained on 880 rice 6mA data and 880 non-6mA data and they did not consider the complex structure information in the sequence such as linkage disequilibrium between nucleotides, thus, there is still some room to improve. Zhou et al. (2018) found 265,290 of rice 6mA data through a variety of experimental methods, such as HPLC-MS/MS, 6mA immunoprecipitation sequencing and Single Molecule Real-Time (SMRT), which enables us to train complex models for 6mA identification. For example, Lv et al. (2019) provided a new random forest model named iDNA6mA-rice based on the reconstructed 154,000 6mA data and 154,000 non-6ma data. iDNA6mA-rice is mainly realized by the random forest algorithm module (RF) based on three feature extraction techniques: K-tuple nucleotide frequency component, mono-nucleotide binary encoding and natural vector. Based on Convolutional Neural Networks (CNN), Yu and Dai (2019) proposed a method named SNNRice6mA to predict the 6mA sites of rice, and showed its advantages over other methods. It is known to us that there is a strong dependence between nucleotides on the sequence, especially on the conserved DNA sequences, thus, the key difficulty in 6mA prediction is to take into consider this dependence structure in statistical modeling. However, as we known, CNN is limited in learning information about long-distance dependence although it has a strong learning ability in the fields of image recognition, voice recognition and agricultural intelligence (10–13). Recurrent Neural Network (RNN) is a special neural network structure, inspired by the fact that human cognition is based on past experience and memory. Different from CNN, RNN not only considers the input of the previous moment, but also can effectively "remember" the previous content. Therefore, RNN has an advantage in analyzing the sequence containing timing information. At present, RNN has been widely used in fields such as natural language processing, image processing, machine translation, speech recognition and bioinformatics (14–16). However, it is difficult to train an RNN due to gradient disappearance or gradient explosion. Long Short-Term Memory (LSTM) and Gate Recurrent Unit (GRU) are proposed to overcome this difficulty, and they are the most commonly used RNNs.

In this study, we introduced a novel deep learning framework named Deep6mA to identify DNA 6mA sites. Deep6mA composed of a CNN and a bidirectional LSTM (BLSTM) module is shown to have a better performance than other methods on 6mA prediction. Interestingly, we find that the motif with the highest frequency of 6mA methylation is concentrated on GAGG among four species: *Rice*, *Arabidopsis thaliana*, *Fragaria vesca*, and *Rosa chinensis*, which means the 6mA methylation has similar patterns across different species. This is further evidenced by the fact that the model trained by rice data has an accuracy over 90% to predict 6mA in the other three species. We may conclude from these results that the sequence prone to DNA 6mA methylation among different species is conservative, and Deep6mA may also be applicable to analyze the 6mA site of other species. More importantly, we found that 6mA is generally enriched in TATA box of the promoter. This may be an important way for 6mA to regulate gene expression.

## MATERIAL AND METHODS

### Benchmark dataset

A benchmark dataset is important to build a reliable prediction model. In this study, for convenience, we use the 6mA-rice-Lv dataset (17,18), including 154,000 positive samples and 154,000 negative samples, to evaluate the proposed method and to compare it with other methods. For each positive sample obtained from GEO, the sequence is 41nt long with the 6mA site locating at the center. For each negative sample collected from NCBI, it’s also a sequence with length 41nt but contains no 6mA modification proved by experiments. In order to demonstrate that 6mA shares the similar patterns across different species and our method can also be used to detect DNA 6mA sites of other species, we also collected DNA 6mA sequences of *Arabidopsis thaliana*, *Fragaria vesca*, and *Rosa chinensis* to show the ability of the trained Deep6mA from rice data on predicting 6mA methylation in these species. The 98483 6mA data of *Arabidopsis thaliana* is obtained from NCBI Gene Expression Omnibus (GEO) with accession number GSE81597, and the database MDR provides 26514 and 14677 6mA data for *Fragaria vesca*, and *Rosa chinensis*, respectively (http://mdr.xieslab.org/).

### Sequence representation

Instead of using manually crafted DNA sequences features, we use the one-hot encoding method to convert the sequence into encoding tensor. Specifically, A, C, G, T, and N are encoded as (1,0,0,0), (0,1,0,0), (0,0,1,0), (0,0,0,1), and (0,0,0,0) respectively. Here the letter ‘N’ represents a non-sequenced nucleotide. Thus, the input DNA sequence is represented as a 4 by 41 encoding matrix, and is viewed as an image which motivates our design of deep learning framework.

### Convolutional neural network and long short term memory network

Convolutional Neural Network (CNN) is widely used in image processing and speech recognition due to its high learning efficiency. The architecture of CNN is analogous to that of the connectivity pattern of neurons in the human brain and was inspired by the organization of the visual cortex. A CNN generally consists of three parts: convolution layers, pooling layers and fully connected layers, which enables it to successfully capture the spatial and temporal dependence in an image. The convolution layer extracts the high-level features such as edges, color and gradient orientation through multiple feature mapping. The resolution of feature mapping is compressed further by a pooling layer to extract dominant features which are rotational and positional invariant, and to decrease the computational power required to process the data. There are two types of Pooling: Max Pooling returning the maximum value from the portion of the image covered by the Kernel and Average Pooling returning the average of all the values from the portion of the image covered by the Kernel. Max Pooling tends to discard the noisy activations and performs de-noising along with dimension reduction; thus, it performs a lot better than Average Pooling. After multiple convolution and pooling processes, one or more fully connected layers are added to learn the non-linear combinations of high-level features from the convolutional layer.

Although CNN is powerful in image processing, however, it does not consider the dependence between inputs, and has a low power in sequence analysis such as natural language processing. Recurrent Neural Network (RNN) is proposed to overcome this shortcoming. As a special type of RNN, Long Short Term Memory Network (LSTM) is not only designed to capture the long dependent information in sequence but also overcome the training difficult of RNN due to gradient explosion and disappearance (19), thus it is the most widely used RNN in real applications. In LSTM, a storage mechanism is used to replace the hidden function in traditional RNN, with a purpose to enhance the learning ability of LSTM for long-distance dependency. Generally, an LSTM unit consists of a storage unit, a forgetting gate, an input gate and an output gate. Bi-directional LSTM (BLSTM), compared with unidirectional LSTM, captures better the information of sequence context.

### The Deep6mA Model

The loci in DNA sequence are known to have a strong linkage disequilibrium, however, it is difficult to take into consider the dependence structure in traditional modeling for predicting 6mA sites. In this section, to fully capture the information in the sequence, we introduce a deep learning network, Deep6mA, which has a CNN to extract high-level features in the sequence and a BLSTM to learn dependence structure along the sequence. Specifically, Deep6mA is consist of five layers of CNN, one BLSTM layer and one fully connected layer. The convolution layer in CNN has 256 filters with filter size 10. The activation function of CNN layers is the exponential linear unit (ReLU).

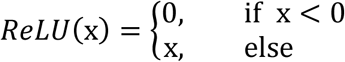

where x is the feature map from the convolution operation. By viewing the input sequence as an image (see Section 2.2), the first convolution layer plays a role as motif detector of the 6mA sites in genome, while the other convolution layers capture higher-level features underlying the sequence. After each convolution layer, a pooling layer with Max Pooling is added to reduce the redundancy of the features from the convolution layer. Then, one BLSTM layer with size 32 is added after CNN to learn the dependence structure in the sequence. The activation function used in this layer is the tanh activation function. A Fully Connected (FC) layer with 32 hidden units is used to learn the non-linear combinations of high-level features from the previous layers. Finally, a sigmoid activation function is used to combine the outputs from the FC layer to make the final decision. Figure 1 shows the flowchart of Deep6mA.

The loss function for training Deep6mA is set as the binary cross-entropy measuring the difference between the target and the predicted output.

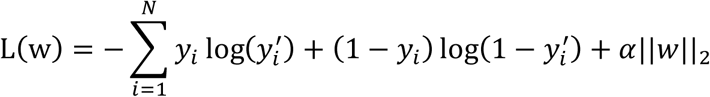

**Figure 1.**
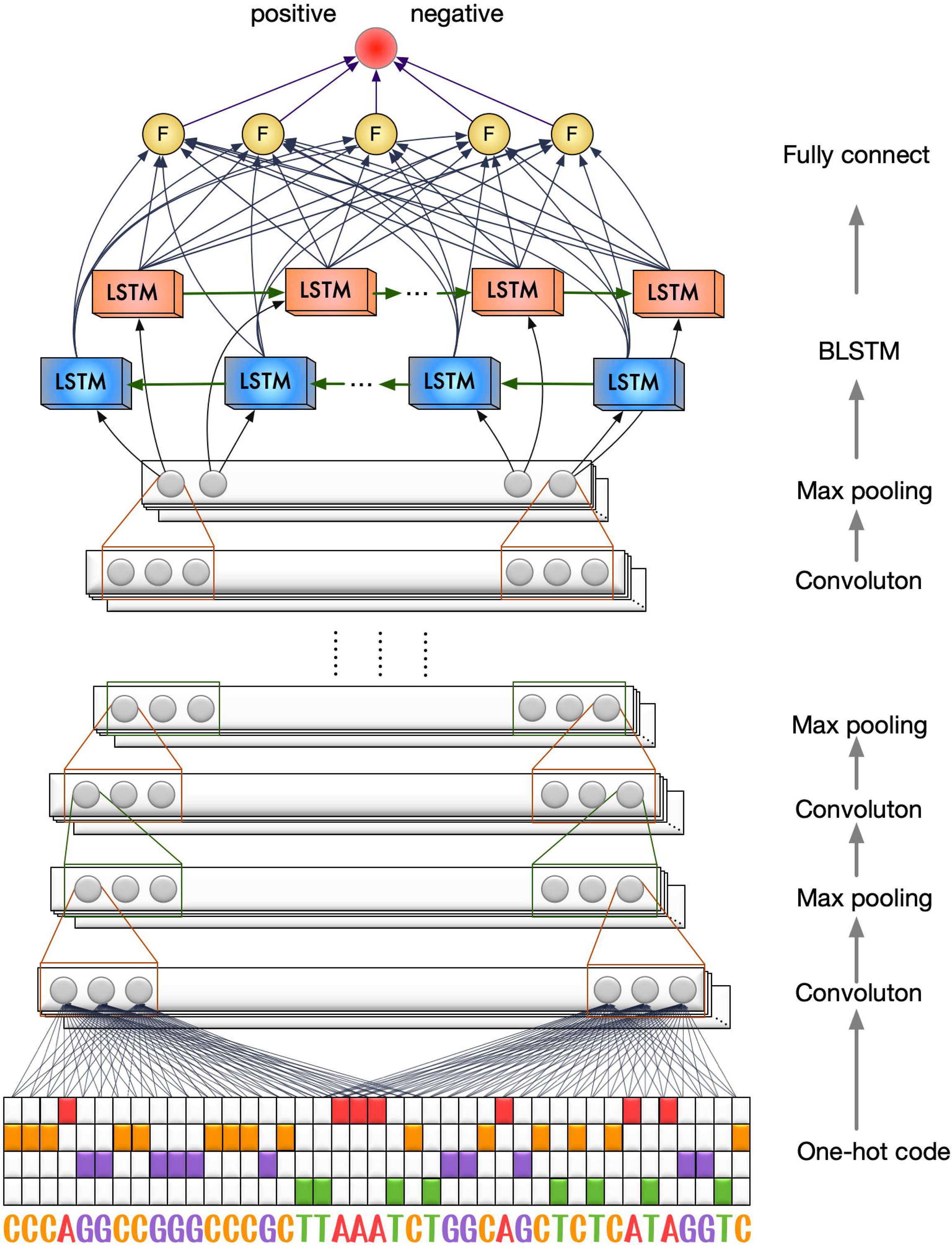
The flowchart of Deep6mA. The structure of Deep6mA consists of five layers of CNN, one layer of BLSTM and one layer of full connection layer.

Where *y*_*i*_ is the true label, *y*′_*i*_ is the corresponding predicted value from Deep6mA, and *α*||*w*||_2_ is a regularization term to avoid overfitting.

Deep6mA is trained by using Adam (20). Batch normalization and dropout (21) are applied after each convolutional procedure to accelerate training and avoid overfitting. When training the model, the dropout rate is set as 0.5, the learning rate is set as 0.01, and the reduced factor is set as 0.5. In addition, the maximum training epoch and batch size is set as 50 and 256, respectively. We take 1/8 of training data, about 10% of the whole dataset, as validation data, and use an early stopping strategy with patience 5, which means the training process will stop when prediction performance did not improve on the validation set. The whole framework is implemented in Pytorch (https://pytorch.org).

### Prediction accuracy assessment

To the best of our knowledge, there are four leading 6mA identification algorithms: i6mA-Pred, SNNRice6mA, iDNA6mA-rice and MM-6mAPred. iDNA6mA-rice is an improved version of i6mA-Pred, and SNNRice6mA is shown to better than iDNA6mA-rice (Yu and Dai, 2019), so we compare Deep6mA with SNNRice6mA and MM-6mAPred. Four metrics, the prediction accuracy (ACC), sensitivity (SN), specificity (SP), and Matthews correlation coefficient (MCC), are used to evaluate the predictive performance of the methods discussed here. Their definitions are given below. A better method should have a higher value on ACC, SN, SP and MCC. The receiver operating characteristic curve (ROC), the area under the curve (11) and precision recall curves (PRC) are used to show the detailed performance of the methods discussed here. The X-axis of the ROC curve is false positive rate (FPR=1-SP), and the Y-axis is true positive rate (TPR=SN). The X-axis of the PRC curve is recall (Recall= SN), and the Y-axis is precision.

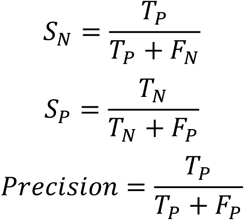

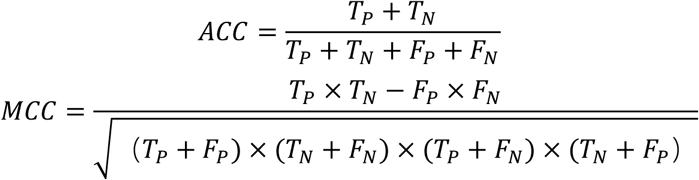

where TP is the number of real 6mA sequences predicted correctly as 6mA methylated, TN is the number of non-6mA sequences correctly predicted as non-6mA methylated, FN is the number of 6mA sequences predicted incorrectly as non-6mA methylated and FP is the number of non-6mA sequences predicted incorrectly as 6mA methylated.

## RESULTS

### Comparing CNN with CNN+LSTM

In this section, we compare the performance of CNN with CNN + LSTM based on the same training data under different settings of CNN. Note that we use the same structure of CNN to compare these two methods. The number and filter of convolution layer in CNN is set to 1and 5 respectively, the corresponding convolution kernel size is set as 32, 64, 128 and 256, respectively, and unit size of LSTM is set to 32. Table 1 is a comparison of the performance of these two methods under different convolution kernels. The result shows that the performance of CNN + LSTM is better than that of CNN, due to the ability of LSTM to learn the dependence structure underlying the sequence.

**Table 1.**
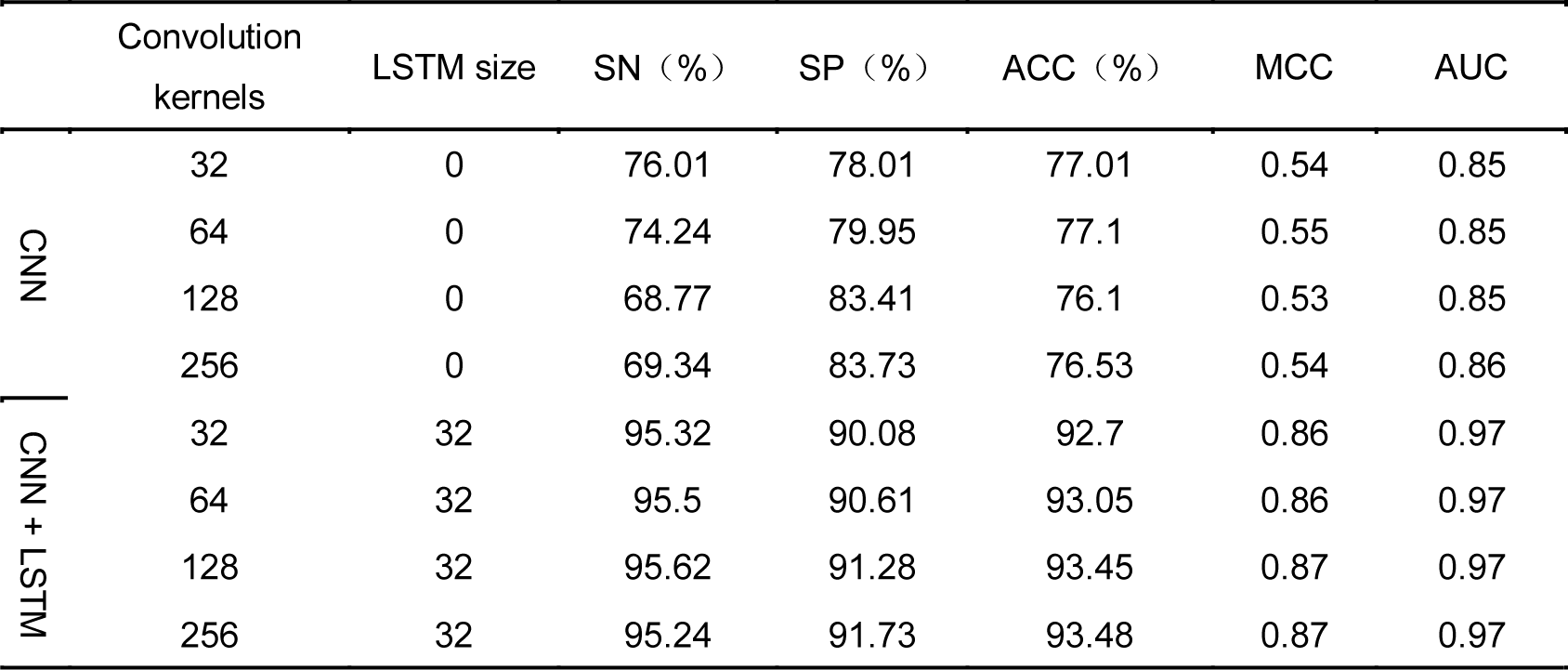
The performance of CNN and CNN + LSTM under different Convolution kernels

|  | Convolution<br>kernels | LSTM size | SN (%) | SP (%) | ACC (%) | MCC | AUC |
| --- | --- | --- | --- | --- | --- | --- | --- |
| CNN | 32 | 0 | 76.01 | 78.01 | 77.01 | 0.54 | 0.85 |
|  | 64 | 0 | 74.24 | 79.95 | 77.1 | 0.55 | 0.85 |
|  | 128 | 0 | 68.77 | 83.41 | 76.1 | 0.53 | 0.85 |
|  | 256 | 0 | 69.34 | 83.73 | 76.53 | 0.54 | 0.86 |
| CNN + LSTM | 32 | 32 | 95.32 | 90.08 | 92.7 | 0.86 | 0.97 |
|  | 64 | 32 | 95.5 | 90.61 | 93.05 | 0.86 | 0.97 |
|  | 128 | 32 | 95.62 | 91.28 | 93.45 | 0.87 | 0.97 |
|  | 256 | 32 | 95.24 | 91.73 | 93.48 | 0.87 | 0.97 |

To better understand the performance of CNN and CNN+LSTM, we show in Figure 2 their ROC curve and PRC curve based on a CNN with 1 convolution layer, 64 convolution kernels, and filter size 10, and a LSTM with 32 hidden units. The AUC of CNN+BLSTM is 0.974, which is 0.124 higher than that of the CNN. Meanwhile, the PR curve also shows that CNN + LSTM is better than CNN.

**Figure 2.**
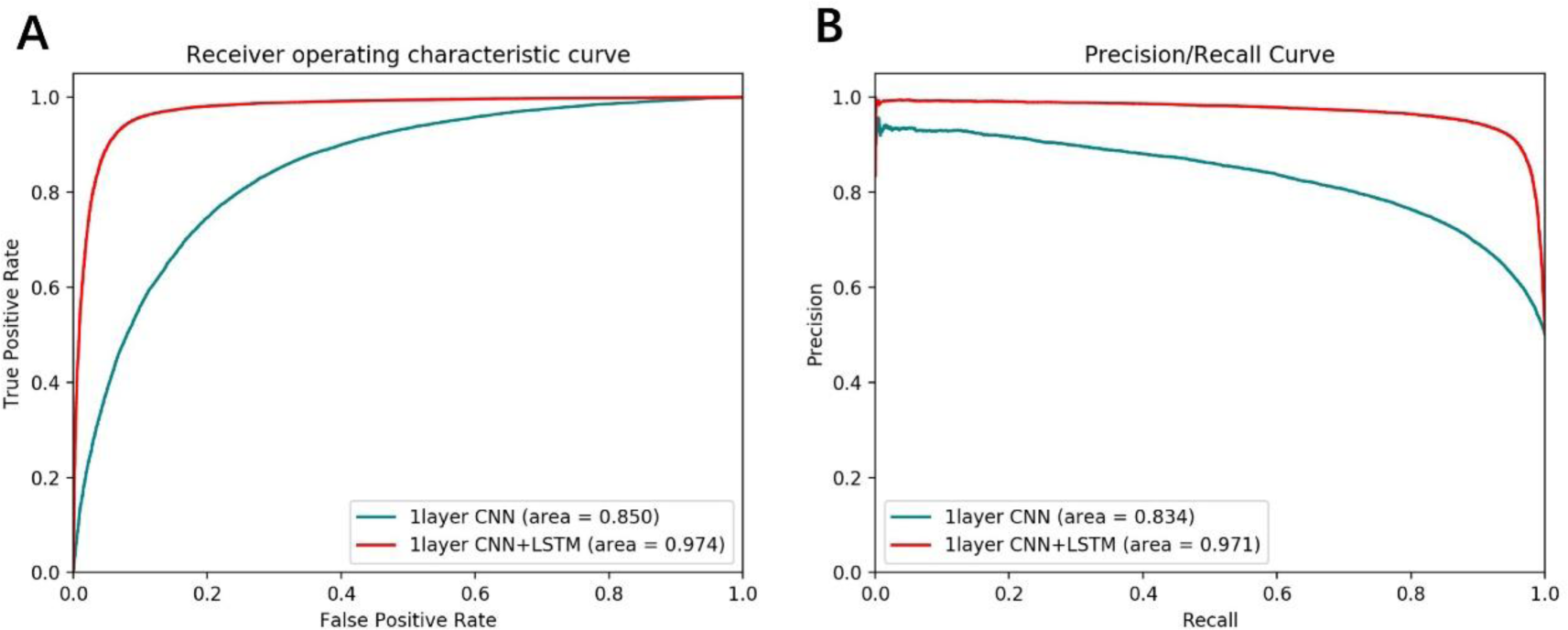
The ROC and PRC curve of CNN and CNN + LSTM based on CNN with 1 convolution layer, 64 convolution kernels, 32 hidden units of LSTM and filter size 10.

### Selecting model parameters of CNN+LSTM

It is known that the performance of CNN+LSTM framework depends on the filter size and the number, convolution kernels of the convolution layer, and number of hidden units in LSTM. To simplify notations, we denote the CNN+LSTM framework with x convolution layer(s), y convolution kernel(s), z filter size and w hidden units as a CNN+LSTM with parameter x-y-z-w. In this section, we select the best CNN+LSTM model from 30 different settings of parameter x, y, z, w by 5-fold cross-validation. Specifically, we take x from {1, 2, 3, 4}, y from {64,256, 512}, w from {16, 32} and fix z at 10. Figure 3 shows the prediction performance of CNN+LSTM model under different settings of parameters. According to these results, the CNN+LSTM with parameter 5-256-10-32 is chosen as our final CNN+LSTEM model.

**Figure 3.**
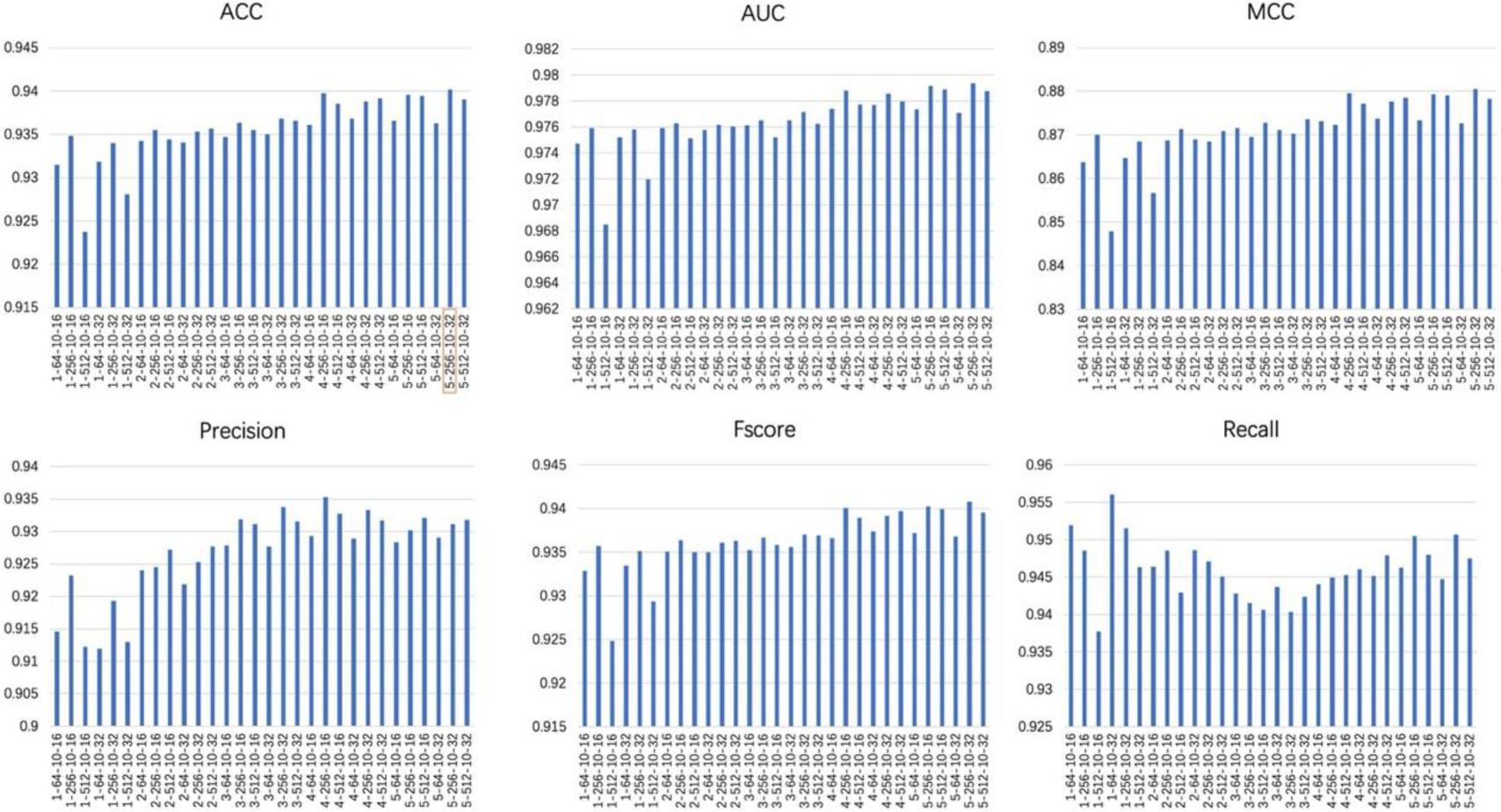
Prediction performance of CNN+LSTM model under different settings of parameters. The CNN+LSTM with parameter 5-256-10-32 is chosen as our final CNN+LSTM model (convolution layer: 5, convolution kernel: 256, filter size: 10, hidden units of LSTM: 32).

### Comparison of motifs across different species

Bailey and Elkan (1995) proposed a MEME algorithm to identify one or more motifs in the sequence based on a binary hybrid model and Expectation-Maximization (EM) algorithm. Here, we use MEME to compare motifs from four species: *Rice*, *Arabidopsis thaliana*, *Fragaria vesca* and *Rosa chinensis*. From Figure 4, we see that GAGG is the most significantly associated motif in these four species. Hence, we infer that DNA 6mA methylation occurs most frequently at GAGG motifs across different species. Xiao et al. (2018) also pointed out that (G/C) AGG (C/T) is the most frequent motif in human genome. All these together suggest that the sequence near the DNA 6mA methylation site may be conserved among different species.

**Figure 4.**
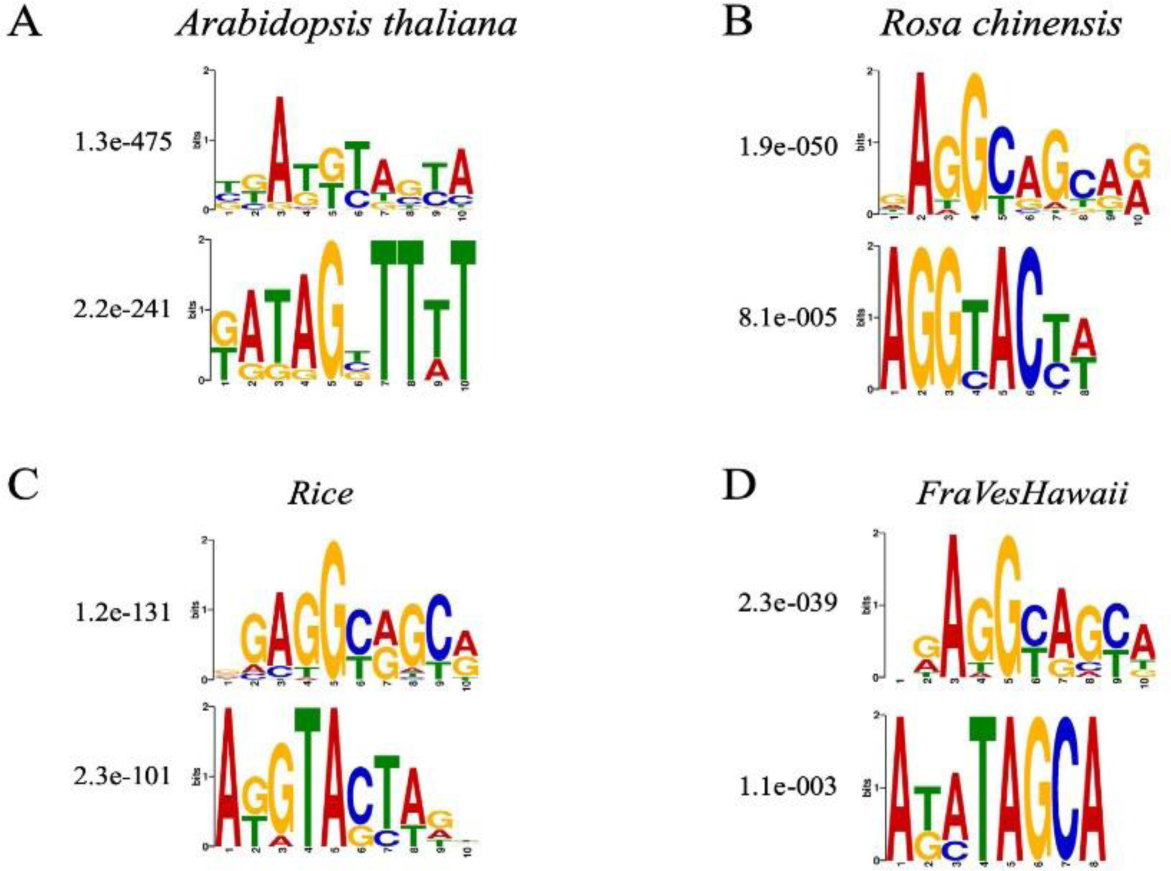
The top two significant motifs in the sequence near the 6mA site of four species (A: Arabidopsis thaliana, B: Rosa chinensis, C: Rice, and D: Fragaria vesca).

### Location features of 6mA sites

It is well known that 5-methylcytosine (5mC) is one of the most important DNA methylation modifications. The results show that the determination of differential methylation region (DMR) is more biologically explanatory and statistically significant than that of the differential methylation sites measured separately (Chen, et al, 2017). Based on this, we expect to know which regions of the genome will be enriched by 6mA methylation. Figure 5A shows the distribution of distance between adjacent 6mA site in 12 chromosomes. It can be seen from Figure 5A that the distribution of 6mA methylation modification on 12 chromosomes is consistent. The mean of the distance between adjacent 6mA sites is greater than 26=64, which means rare 6mA sites occur in a continuous region like 5mC sites to form a DMR. To explore further the location features of 6mA sites, we look inside the subsequences of length 30nt with more than five 6mA sites, and find that almost all of the 6mA sites in these subsequences are located in the TATA box of the promoter (see Figure 5B for example, more details are given in Supplementary Material). This implies that 6mA is generally enriched in TATA box which is a very important functional component of the promoter. The transcription process will not start until RNA polymerase binds tightly on TATA box. Therefore, the enrichment of 6mA methylation on TATA box may directly affect the transcription and expression of downstream genes. This may be an important regulatory function of 6mA methylation modification.

**Figure 5.**
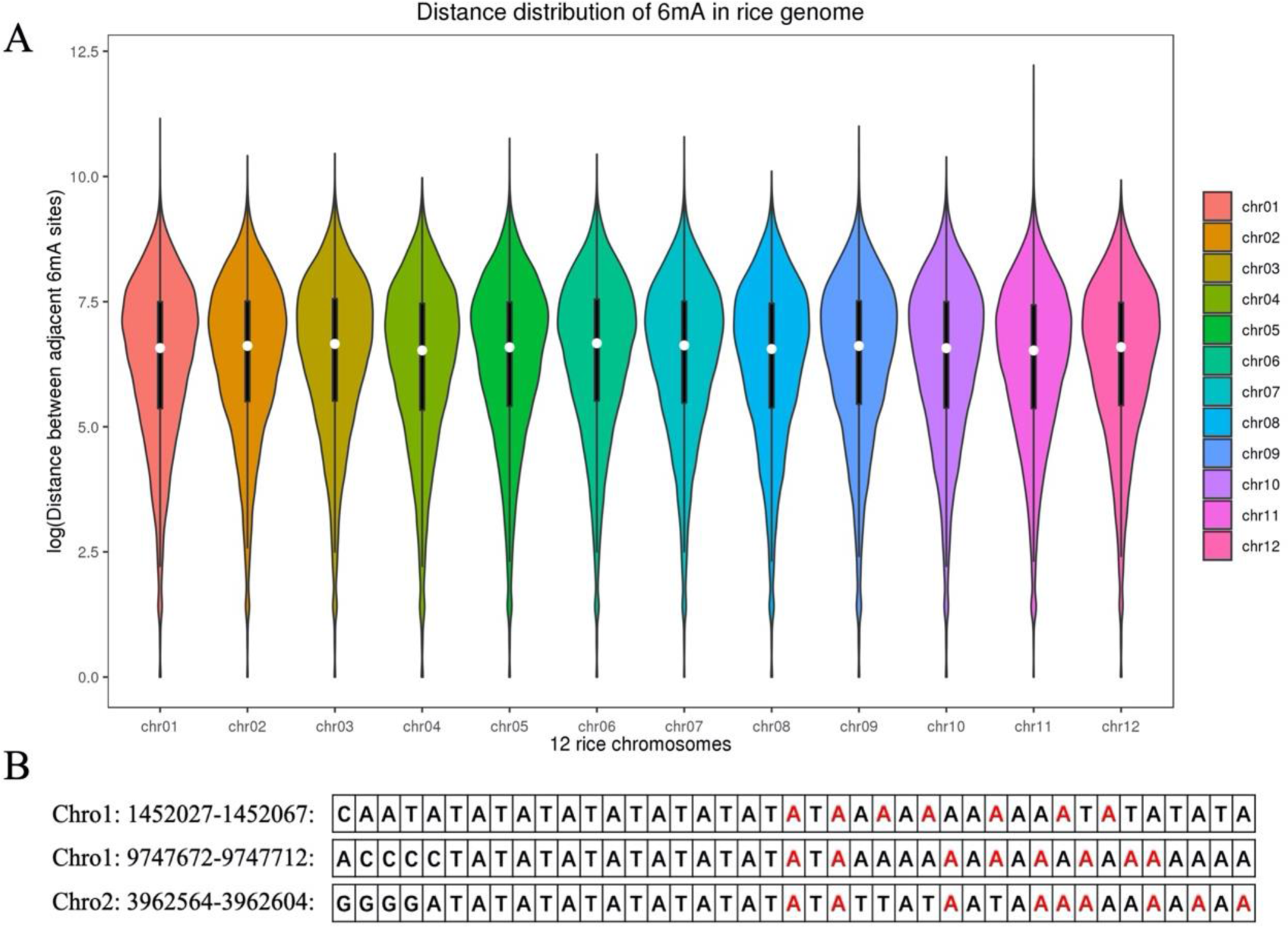
Location features of 6mA sites in the sequence. (A) The distribution of 6mA methylation modification on 12 chromosomes. The X axis and Y axis represent the 12 chromosomes and the logarithm of distance between adjacent 6mA site. (B) Three examples of sequences containing TATA box enriched with 6mA methylation (colored in red).

### Comparison with other leading methods

We use 5-fold cross-validation to evaluate the performance of Deep6mA, SNNRice6mA and MM-6mAPred based on 6mA-rice-Lv dataset (see Section 2.1). The settings of the iDNA6mA-rice and SNNRice6mA follow that given in the corresponding original paper. Specifically, the number of trees in forest algorithm (RF) in iDNA6mA-rice is set to 100 with the seed of 1. For SNNRice6mA, the number of convolution layer and convolution kernel is set to 1 and 4, respectively, and the filter size and fully connect layer size of SNNRice6mA is set to 3 and 64, respectively. Deep6mA is the best one among these methods in terms of SN, SP, ACC and MCC as shown in Table 2 with SN, SP, ACC and MCC as 95.06%, 92.96%, 94.01% and 0.88 respectively. The better performance of Deep6mA is mainly due to the ability of BLSTM to learn the dependence structure between distant nucleotides.

**Table 2.** Comparison of Deep6mA, SNNRice6mA and MM-6mAPred based on 6Ma-rice-Lv dataset.

| Method | SN (%) | SN (%) | ACC (%) | MCC | AUC |
| --- | --- | --- | --- | --- | --- |
| Deep6mA | 95.06 | 92.96 | 94.01 | 0.88 | 0.98 |
| SNNRice6mA-large | 94.33 | 89.75 | 92.04 | 0.84 | 0.97 |
| MM-6mAPred | 93.47 | 89.51 | 91.49 | 0.83 | 0.96 |

In addition, the ROC curve and PR curve of Deep6mA, SNNRice6mA and MM-6mAPred are shown in Figure 6. The area under curve of Deep6mA is 0.979, which is higher than that from other two methods. All these results show that our method Deep6mA is the best one among these methods.

**Figure 6.**
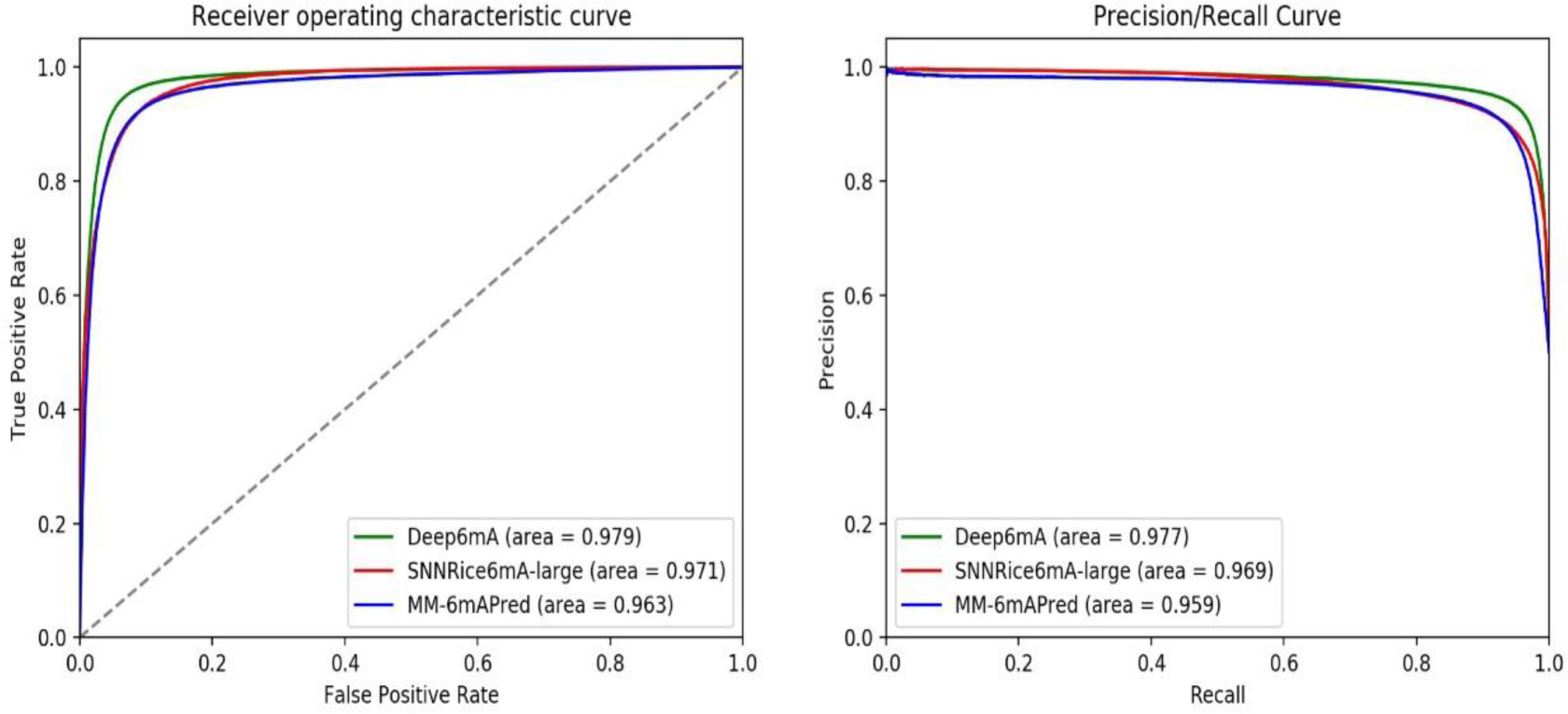
The ROC curves (A) and PRC curves (B) of Deep6mA based on 6mA-rice-Lv dataset.

### Validation on other three species

Results in Section 3.3 show that 6mA is conservative among different species, which suggest that Deep6mA trained on rice data is applicable to predict the 6mA sites of other species. In the following, we try to validate this principle by applying the trained Deep6mA to the 6mA data of other three species: *Arabidopsis thaliana* with sample size 98483, *Fragaria vesca* with sample size 26514 and *Rosa chinensis* with sample size 14677 (see Section 2.1 for details). The prediction results on these three test datasets are listed in Table 3. We found that Deep6mA trained with rice 6mA data predicts with high accuracy the 6mA sites in other three species. The prediction performance of our method is better than SNNRice6mA and MM-6mAPred. This makes deep6mA have great potential in predicting 6mA sites in other species’ genomes.

**Table 3.**
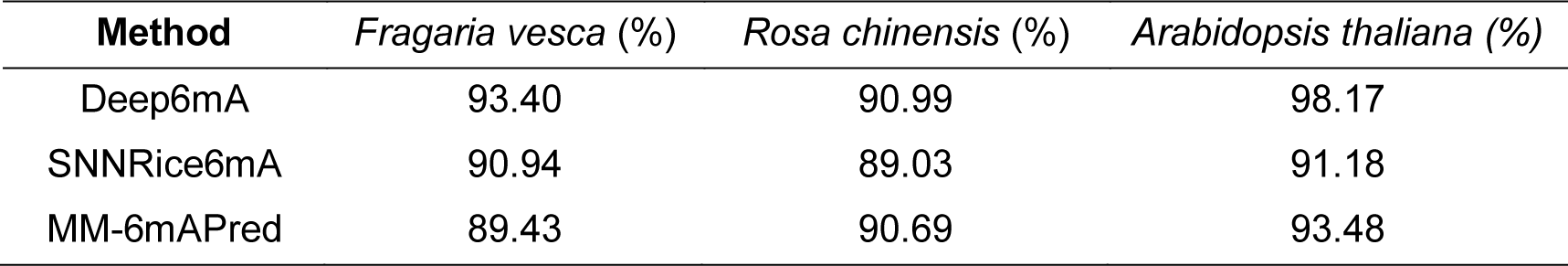
Prediction accuracy of Deep6mA, SNNRice6mA and MM-6mAPred trained with rice data on Arabidopsis thaliana, Fragaria vesca and Rosa chinensis.

## DISCUSSION

In this study, we propose a deep learning framework named Deep6mA by integrating CNN and LSTM to efficiently predict DNA 6mA sites. Deep6mA uses a CNN layer to extract DNA sequence characterization, and then spreads it into a BLSTM layer to capture context dependency information of 6mA sites. Finally, these features are transferred to a fully connected layer to determine whether the site is a 6mA site. The experimental results show that Deep6mA can predict the 6mA site of rice with high accuracy. Importantly, we found that most of the 6mA methylation modifications in different species are more likely to occur on GAGG motifs. This shows that DNA sub-sequences containing 6mA sites among species have certain conservation. Maybe this is the reason why Deep6mA model trained with rice data can accurately predict 6mA sites in other three species. However, there are some inadequacies in this study, such as the selection of sequence length. In theory, the longer the sequence, the more information it provides. All previous studies on 6mA site recognition are based on the sequence with a length of 41nt. It is not necessary to learn the complete sequence information by only training the model with those short sequences. Besides, due to the relative complexity of the calculation time, the framework and parameter design of deep6mA may only achieve a local optimum. What’s more, why is the 6mA site enriched in the TATA box of the promoter, and whether this enrichment has a regulatory effect on the expression of downstream genes. For the ongoing work, we will carry out further research on these issues

## DATA AVAILABILITY

Source code, all training data and testing data of Deep6mA are found at https://github.com/Marscolono/Deep6mA

## SUPPLEMENTARY DATA

Supplementary Data are available at NAR online.

## ACKNOWLEDGEMENT

We would like to thank Prof. Xiaodan Fan from The Chinese University of Hong Kong for valuable discussions.

